# Maltodextrin Transport in the Extremely Thermophilic, Lignocellulose Degrading Bacterium *Anaerocellum bescii (f. Caldicellulosiruptor bescii)*

**DOI:** 10.1101/2024.09.14.613025

**Authors:** Hansen Tjo, Virginia Jiang, Jerelle A. Joseph, Jonathan M. Conway

**Affiliations:** Department of Chemical and Biological Engineering, Princeton University, Princeton, NJ 08544, USA; Omenn-Darling Bioengineering Institute, Princeton University, Princeton, NJ 08544, USA; Andlinger Center for Energy and the Environment, Princeton University, Princeton, NJ 08544, USA; High Meadows Environmental Institute, Princeton University, Princeton, NJ 08544, USA

**Author notes:** Corresponding Author: Jonathan M. Conway.

**Keywords:** thermophile, biophysics, lignocellulose, ABC sugar transporter, substrate-binding protein, *Caldicellulosiruptor*, *Anaerocellum bescii*

## Abstract

Sugar transport into microbial cells is a critical, yet understudied step in the conversion of lignocellulosic biomass to metabolic products. *Anaerocellum bescii* (formerly *Caldicellulosiruptor bescii*) is an extremely thermophilic, anaerobic bacterium that readily degrades the cellulose and hemicellulose components of lignocellulosic biomass into a diversity of oligosaccharide substrates. Despite significant understanding of how this microorganism degrades lignocellulose, the mechanisms underlying its highly efficient transport of the resulting oligosaccharides into the cell are comparatively underexplored. Here, we identify and characterize the ATP-Binding Cassette (ABC) transporters in *A. bescii* governing maltodextrin transport. Utilizing past transcriptomic studies on *Anaerocellum* and *Caldicellulosiruptor* species, we identify two maltodextrin transporters in *A. bescii* and express and purify their substrate-binding proteins (Athe_2310 and Athe_2574) for characterization. Using differential scanning calorimetry and isothermal titration calorimetry, we show that Athe_2310 strongly interacts with shorter maltodextrins such as maltose and trehalose with dissociation constants in the micromolar range, while Athe_2574 binds longer maltodextrins, with dissociation constants in the sub-micro molar range. Using a sequence-structure-function comparison approach combined with molecular modeling we provide context for the specificity of each of these substrate-binding proteins. We propose that *A. bescii* utilizes orthogonal ABC transporters to uptake malto-oligosaccharides of different lengths to maximize transport efficiency.

**Importance:** Here, we reveal the biophysical and structural basis for oligosaccharide transport by two maltodextrin ABC transporters in *A. bescii*. This is the first biophysical characterization of carbohydrate uptake in this organism and establishes a workflow for characterizing other oligosaccharide transporters in *A. bescii* and similar lignocellulosic thermophiles of interest for lignocellulosic bioprocessing. By deciphering the mechanisms underlying high affinity sugar uptake in *A. bescii*, we shed light on an underexplored step between extracellular lignocellulose degradation and intracellular conversion of sugars to metabolic products. This understanding will expand opportunities for harnessing sugar transport in thermophiles to reshape lignocellulose bioprocessing as part of a renewable bioeconomy.

## Introduction

Using lignocellulosic biomass as a source of renewable fuels and chemicals is critical to addressing today’s energy and environmental challenges (1–3). Though lignocellulosic biomass conversion is constrained by its physical and chemical recalcitrance to degradation, nature offers a promising solution: cellulolytic and hemicellulolytic thermophiles (4,5). These microbes have evolved to thrive in sugar-limiting environments by efficiently consuming a wide-range of monosaccharide and oligosaccharide sugars (6–8). This adaptation makes them especially valuable in biotechnology contexts, particularly for converting lignocellulosic biomass into high-value fuel and chemical commodities, e.g., ethanol or acetone (7,9,10).

*Anaerocellum bescii*, an extremely thermophilic bacterium (optimum growth temperature of 78°C), holds exceptional promise as a metabolic engineering workhorse due to its highly effective carbohydrate-active enzymes (CAZymes), absence of carbon catabolite repression enabling simultaneous utilization of a broad range of substrates, and availability of a genetic system for metabolic engineering (11–13). These strengths make

*A. bescii* especially promising for consolidated bioprocessing of lignocellulosic biomass (9,14,15). Multiple studies have shown how *A. bescii* can be genetically engineered to produce such valuable chemical commodities as acetone, ethanol, and acetoin from lignocellulose at industrially-relevant yields (9,16,17). Yet, this substantial breadth of research into engineered metabolism in *A. bescii* overshadows our limited understanding on how it utilizes diverse lignocellulosic substrates (18,19). To fully harness its potential as a robust metabolic workhorse, a more fulsome understanding of the mechanisms governing sugar transport in *A. bescii* is vital.

Transcriptomics have shed light on sugar transport and utilization in the *Caldicellulosiruptor* genus (recently split into two genera: *Caldicellulosiruptor* and *Anaerocellum*) (13,18,19). VanFossen *et al*. (2009) examined *C. saccharolyticus,* proposing putative substrates for all twenty-four of its ATP-Binding Cassette (ABC) sugar transporters (18). These results were expanded by a metabolic model from Rodionov *et al*. (2021) on genetically similar *Anaerocellum bescii*, with updated genomic annotation on a broader range of carbohydrate utilization genes spanning transcriptional regulators, as well as intracellular and extracellular CAZymes (19). New predictions of each ABC transporter’s substrate specificity were also made, though to date have not been experimentally verified.

High-affinity ATP-Binding Cassette (ABC) transporters mediate sugar transport in lignocellulolytic thermophiles such as *A. bescii* (8,19–22). These ABC transporters typically comprise an extracellular substrate-binding protein, transmembrane domain-containing proteins, and an intracellular ATPase (23,24). ABC transporters are also divided into two families: Carbohydrate-Uptake-Transporter Family 1 (CUT1) and Carbohydrate-Uptake-Transporter Family 2 (CUT2). CUT1 transporters include a heterodimeric transmembrane pore made of two different proteins, whereas CUT2 transporters utilize a homodimeric transmembrane barrel (23,24). CUT1 transporters also primarily uptake larger oligosaccharides, while CUT2 transporters preferentially uptake disaccharides and monosaccharides (23,24). For both transporter types, hydrolysis of two ATP molecules by the intracellular ATPase powers the translocation of one sugar molecule.

Because the substrate-binding domain is responsible for binding and delivering sugars in the extracellular space to the transmembrane entryway, it largely dictates an ABC transporter’s substrate specificity (22,25,26). As thermophiles like *A. bescii* are natively found in nutrient limiting environments, substrate-binding proteins crucially enhance sugar uptake efficiency by increasing the frequency of substrate-transporter interactions (27). Substrate-binding proteins typically contain two domains connected by a hinge region. The presence of the appropriate ligand drives flexion about the hinge region and movement of one domain towards the other to capture the ligand in the binding pocket. With a bound substrate, the substrate-binding protein adopts a closed – and more thermally stable – conformation (22,25,28). This so-called “Venus Fly-trap” mechanism has been studied extensively, with the degree of protein closure dependent on binding affinity to the substrate (25,29). Substrate-binding proteins that bind malto-oligosaccharides (commonly referred to as MBPs), possess multiple subsites within their binding pocket for ligand coordination (30–32). Each glucosyl residue of a maltodextrin is positioned in a sub-site, stabilized by a combination of hydrogen bonds and steric interactions. The non-reducing glucosyl ring of maltodextrin ligands typically occupies the most deeply buried sub-site, with its C6 hydroxyl group forming a “capping” hydrogen bond, constraining the possible conformations between ligand and protein (30). Thus, substrate specificity depends on both the availability and residue composition of individual subsites in the binding pocket.

ABC transporters for the uptake of maltodextrins have represented a classical model with which to understand sugar transport in bacteria (21,22,33,34). The ABC transporter for maltodextrin transport in *E. coli,* organized as *malEFGK* operons, is a prime historical example (35–37). These transporter Open Reading Frames (ORFs) typically, though not exclusively, encode for the maltodextrin-binding protein (MalE, also referred to as MBP), two transmembrane domains (MalF and MalG), and the ATPase subunit (MalK) (38–40). Recently, a wealth of structural and biophysical information on maltodextrin-binding proteins in several notable thermophilic bacteria and archaea have been generated: *Thermococcus litoralis, Thermotoga maritima, Thermus thermophilus,* and *Pyroccocus furiosus* (20,21,32,33,39). While these thermophilic orthologs share similarities in structural folds and an affinity for ɑ-linked glucans, their precise substrate specificities differ. For instance, *Thermotoga maritima* and *P. furiosus* possess maltodextrin-binding proteins that preferentially bind longer oligosaccharides such as maltotriose (21,32). Moreover, there has been limited characterization of ABC sugar transport in extreme thermophiles that have gained recent prominence for their consolidated bioprocessing capabilities, particularly those of the *Anaerocellum* and *Caldicellulosiruptor* genera.

Here, we present the first ever biophysical analysis of the maltose/trehalose and malto-oligosaccharide ABC transporters, Athe_2310 and Athe_2574, from the extremely thermophilic *Anaerocellum* or *Caldicellulosiruptor* genera. Each transporter’s extracellular substrate-binding proteins were heterologously produced in *E. coli* for *in vitro* study. We employ both differential scanning calorimetry (DSC) and isothermal titration calorimetry (ITC) to identify the substrate specificity of these transporters, showing Athe_2310 binds maltose and trehalose, while Athe_2574 binds larger malto-oligosaccharides. Using crystal structures of homologous proteins, we elucidate specific binding pocket residues in these proteins that are highly conserved for maltodextrin binding. Finally, we develop a tailored-to-design ligand docking model that builds on predicted structures of Athe_2310 and Athe_2574 from AlphaFold2 to more accurately model maltodextrin-binding. By experimentally determining the substrate specificity of Athe_2310 and Athe_2574, and elucidating the structural context that underpins their differences, we show that *A. bescii* relies on a network of high-affinity ABC transporters with distinct substrate specificity for sugar uptake.

## Results

### Identification of the Maltodextrin ABC Transport System in *A. bescii*

*A. bescii* contains two loci encoding ABC transporters (Athe_2308 - Athe_2310 and Athe_2574 - Athe_2578) predicted to mediate maltodextrin transport based on transcriptomic work by VanFossen *et al*. (2009) in *Caldicellulosiruptor saccharolyticus* and Rodionov *et al*. (2021) in *A. bescii* (18,19). Originally, VanFossen *et al*. (2009) predicted that Csac_2493 (homologous to Athe_2310) binds xylose, glucose, and fructose, while Csac_0431 (homologous to Athe_2574) binds maltodextrins (18). These predictions were based on differential expression of ORFs when *C. saccharolyticus* was grown on pentose versus hexose monosaccharides, a homology search in *T. maritima,* and the specificity of any genomically co-localized glycoside hydrolases. More recently, Rodionov *et al*. (2021) predicted the presence of two putative ABC transporters for maltodextrin transport encoded by the *malEFG* (containing Athe_2574) and *malEFG2* (containing Athe_2310) operons. This was based on operon identification through homology searches and genome context identification, followed by transcriptome sequencing analysis on *A. bescii* grown on separate hexose and pentose sugar sources – revealing co-upregulation of the putative *malEFG, malEFG2* and starch degradation operons *(pulA/amyX)* on cellobiose and Avicel (19). However, all the above transporter substrate predictions based on transcriptomics have not been further validated.

Inspection of genomic context suggests that Athe_2574 is indeed a putative maltodextrin transporter (Figure 1). Substrate binding domain Athe_2574 has neighboring genes associated with ɑ-glucan metabolism (such as ɑ-glucan phosphorylase Athe_2576 and ɑ-amylase Athe_2579) and the transmembrane proteins Athe_2577-78. The Athe_2308 - Athe_2310 locus (Figure 1), however, lacks any proximal genes involved in polysaccharide degradation or metabolism that would have suggested the substrate preference for this transporter. Both transporters contain two transmembrane domain ORFs, (Athe_2577-78) and (Athe_2309-10), implying that both ABC transporters are CUT1 family members, which typically have a preference for oligosaccharides (23). Neither locus contains a MalK or another ATPase. It was recently discovered that all CUT1 transporters in *A. bescii* are powered by a promiscuous ATPase, encoded by *msmK* (Athe_1803) (19). Presumably both putative maltodextrin transporters utilize the MsmK ATPase.

**Figure 1:**
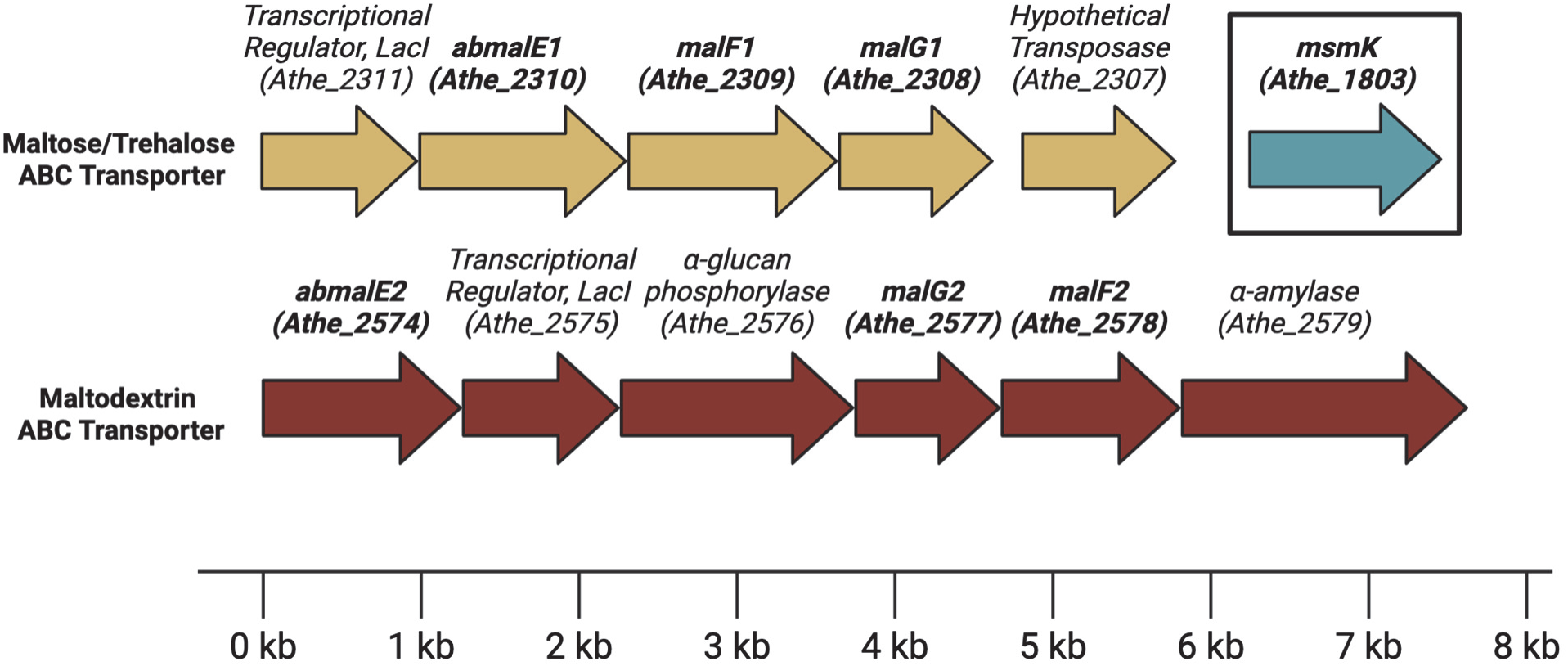
Genomic organization of maltodextrin ATP-binding cassette (ABC) transporters and neighboring genes associated with amylose metabolism in *Anaerocellum bescii*.

### Sequence and Structural Comparison to Thermophilic Orthologs

As there are no resolved liganded structures of Athe_2310 and Athe_2574, crystal structures of holo orthologs in Protein Data Bank (PDB) and predictive structures from AlphaFold2 can provide useful insights into each protein’s respective general fold as well as ligand binding pocket (41). Athe_2310 is closest in protein sequence to TmMBP3 (PDB: 6DTQ, *Thermotoga maritima*), TMBP (PDB: 1EU8, *Thermococcus litoralis*), and TtMBP (PDB: 6J9W, *Thermus thermophilus*), with a coverage of over 90% for all three orthologs and an amino acid sequence identity ranging from 31.7% to 37.4% (Table 1). Athe_2574 is most similar to PfuMBP (PDB: 1ELJ, *Pyrococcus furiosus*), TmMBP1 (PDB: 6DTU, *Thermotoga maritima*), and TmMBP2 (PDB: 6DTS, *Thermotoga maritima*), with an amino acid sequence identity ranging from 31.1% to 38.5% with over 92% coverage (Table 1). Closer inspection of these groups of orthologs reveal that they bind ɑ-glucans of similar size. Proteins TmMBP3, TMBP, and TtMBP all bind ɑ-linked disaccharides, while TmMBP1, TmMBP2, and PfuMBP all bind larger maltodextrins such as maltotriose (G3) and maltotetraose (G4). These comparisons suggest that Athe_2310 may preferentially bind disaccharides, while Athe_2574 preferentially binds larger maltodextrins. Table 1 contains a summary of these holo orthologs and their similarity by BLASTp to Athe_2310 and Athe_2574.

**Table 1:**
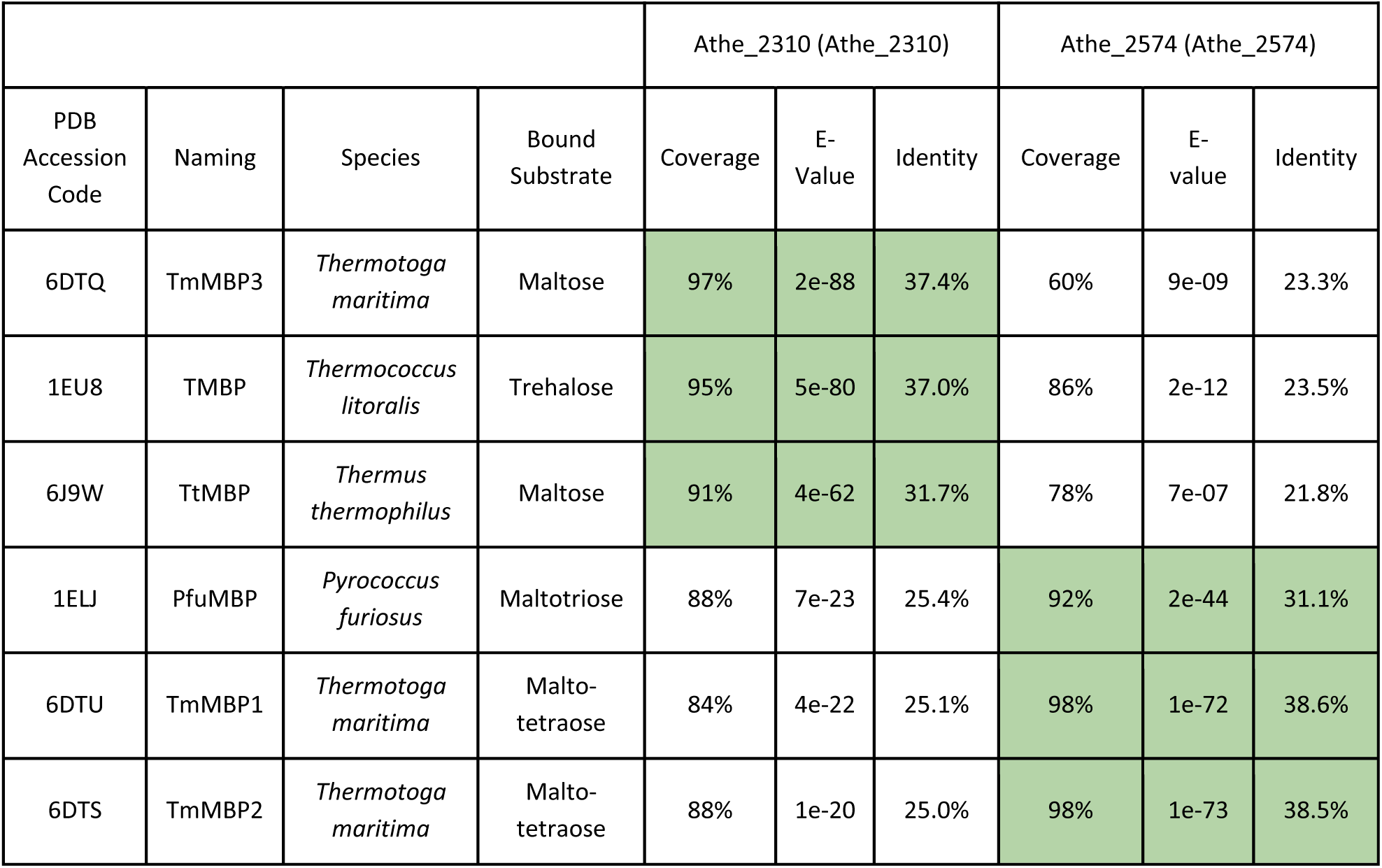
BLASTp local alignment of Athe_2310 and Athe_2574 to a panel of structurally resolved and liganded thermophilic orthologs indicate divergent sequence similarities and substrate preferences. E-values represent the probability of finding a hit of similar quality by random chance.

Like their thermophilic orthologs, Athe_2310 and Athe_2574 are structured as two protein domains connected by a hinge region comprising an ɑ-helix and anti-parallel β-strands. In both MBPs, their ɑ-helices and β-strands are distributed across their two domains. Their respective ligand binding sites, further visualized in Figures 2a and 3a, are in a furrow that lies opposite the hinge region. From our multiple-sequence alignment on PROMALS3D, we mapped the full list of Athe_2310 and Athe_2574 residues that align with fully conserved binding pocket residues across their respective orthologous subsets; we predict these residues are similarly important in Athe_2310 and Athe_2574 for maltodextrin coordination. In both Figures 2 and 3, we classified certain residues as being fully conserved (colored green), or partially conserved (colored orange), with respect to their orthologous subsets. Finally, Athe_2310 and Athe_2574 also differ in solvent accessibility and electrostatic potential. The pocket volume of Athe_2310 is 109 Å^3^, much smaller than Athe_2574, which has a pocket volume of 396 Å^3^.

**Figure 2:**
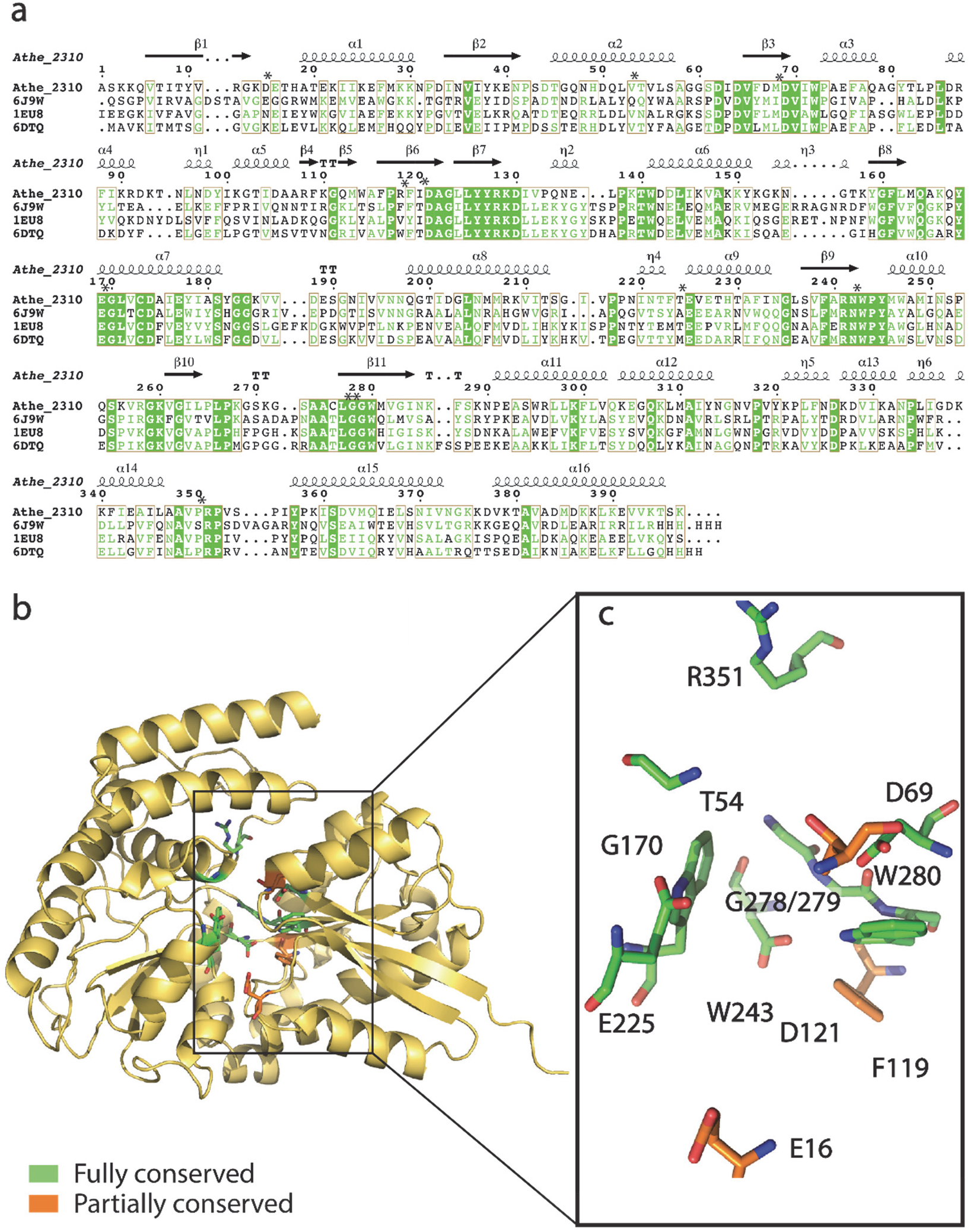
a) Structure-based amino acid sequence alignment of Athe_2310 with its thermophilic holo orthologs: TtMBP (PDB: 6J9W), TMBP (PDB: 1EU8), TmMBP3 (PDB: 6DTQ). Residues shaded in green indicate full residue conservation in Athe_2310 and all orthologs. Residues in green indicate chemically similar residues. Boxes are drawn wherever aligned residues are chemically similar for at least three out of proteins (sequence consensus with a threshold of >70%). Binding pocket residues are marked with an asterisk. b) PyMol visualization of the AlphaFold2 structure of Athe_2310. Binding pocket residues in Athe_2310 are marked as either fully conserved (highlighted in green) or partially conserved (highlighted in orange) compared to its orthologs 6J9W, 1EU8, 6DTQ. c) Visualization and labeling of both fully and partially conserved residues in the Athe_2310 binding pocket.

**Figure 3:**
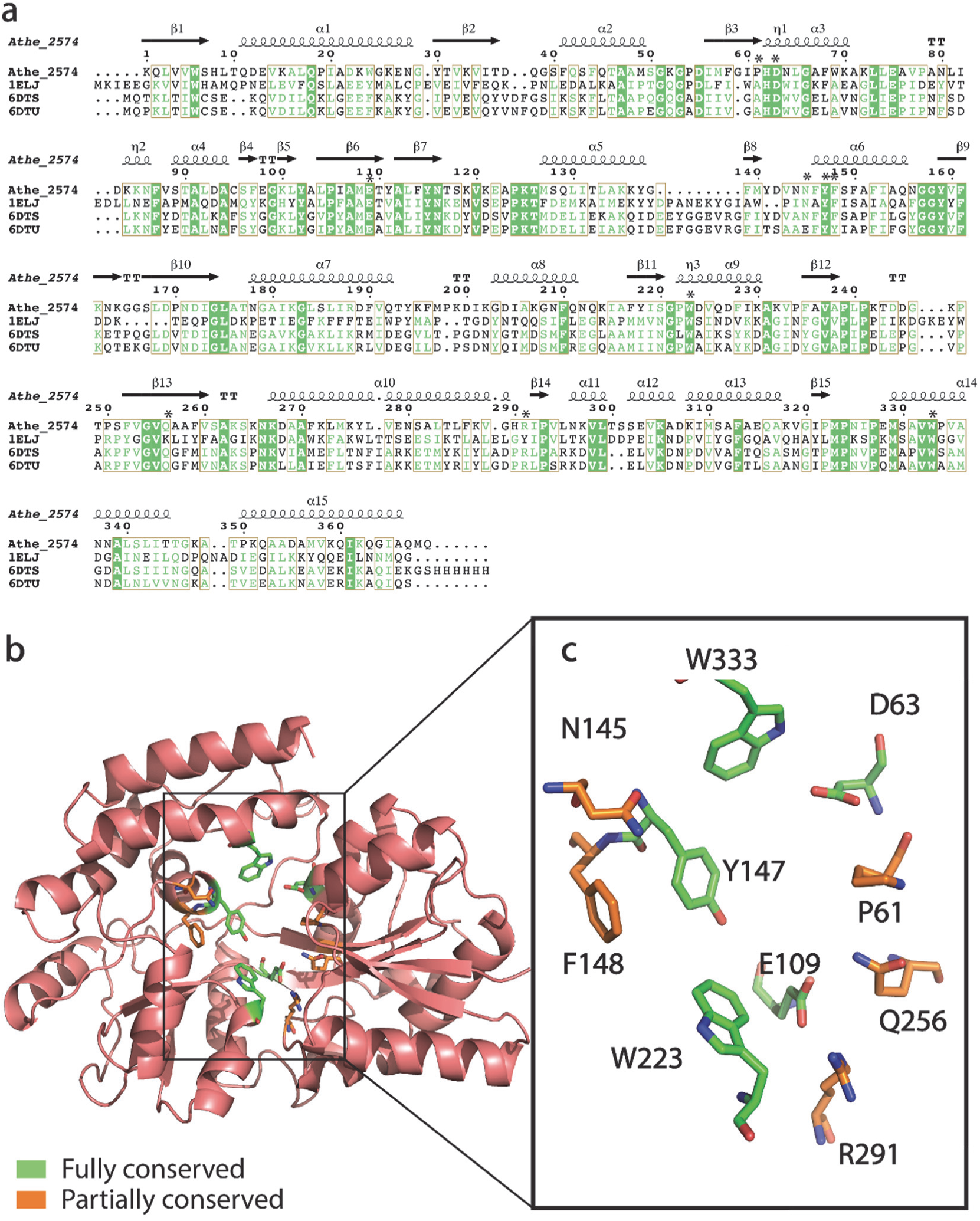
a) Structure-based amino acid sequence alignment of Athe_2574 with its thermophilic holo orthologs: PfuMBP (PDB: 1ELJ), TmMBP2 (PDB: 6DTS), TmMBP1 (PDB: 6DTU). Residues shaded in green indicate full residue conservation in Athe_2310 and all orthologs. Residues in green indicate chemically similar residues. Boxes are drawn wherever aligned residues are chemically similar for at least three out of proteins (sequence consensus with a threshold of >70%). Binding pocket residues are marked with an asterisk. b) PyMol visualization of the AlphaFold2 structure to Athe_2574. Binding pocket residues in Athe_2574 are marked as either fully conserved (highlighted in green) or partially conserved (highlighted in orange) as compared to its orthologs 1ELJ, 6DTS, 6DTQ. c) Visualization and labeling of both fully and partially conserved residues in the Athe_2574 binding pocket.

The binding pocket of Athe_2310 is closest in sequence composition and subsite packing to orthologs TMBP, TmMBP3, and TtMBP (Figure 2). In Athe_2310, F119 and W243 appear to be critical in fostering pi–pi stacking interactions with bound carbohydrates, while G278 and Gly279 are likely implicated in hydrophobic interactions (42–44). As is the case in TtMBP, it is also possible D69, W243, and R351 coordinate both movement of the N-terminal subunit upon ligand recognition and stabilizing interactions in the binding pocket (20). Binding pocket residues in Athe_2310 are also more deeply conserved than in Athe_2574. While Athe_2574 exhibits comparatively fewer conserved binding pocket residues (Figure 3), phenyl residues including Y147, W223, and W333 suggest a similar pi–pi stacking mechanism with hexose rings (42). D63 and Y147 may be involved in forming hydrogen bonds with glucose oxygen atoms, as is the case between conserved aspartic acid and tyrosine residues in PfuMBP (D68 and Y161) with maltotriose (21).

### Biophysical Determination of Substrate Specificity

In DSC, proteins are heated at a constant rate to induce thermal denaturation. This can be measured by an increase in the protein’s specific heat capacity *C*_p_(*T*) as a function of temperature. The melting temperature *T*_m_ is defined as the temperature at which the protein attains its maximum heat capacity and becomes fully denatured (45). However, the presence of their cognate ligands can induce substrate-binding proteins to adopt a more thermally stable closed conformation, resulting in an increase in *T*_m_ (46). This *ΔT*_m_ = |*T*_m,holo_ - *T*_m,apo_| difference can be used to determine differential substrate specificity, as preferred ligands that are more tightly bound will yield larger *ΔT*_m_ values. DSC results show that Athe_2310 exhibits the largest melting temperature shift for maltose substrate, with *ΔT*_m_ = 9.06 °C (Figure 4a). However, other short ɑ-glucans trehalose (*ΔT*_m_ = 7.23 °C) and maltotriose with (*ΔT*_m_ = 6.63 °C) also yield substantial melting temperature differences. Melting temperature differences are subtler with longer oligosaccharides maltotetraose (*ΔT*_m_ = 1.57 °C), maltopentaose (*ΔT*_m_ = 0.95 °C), and maltoheptaose (*ΔT*_m_ = 0.26 °C) (Figure 4a). However, as binding of maltose to *E. coli* MBP is known to increase protein melting temperature by approximately 8 °C, depending on pH, it is unlikely binding is taking place between Athe_2310 and longer maltodextrins (G4–G7) due to their much smaller *ΔT*_m_ values in comparison (46,47). The presence of a single peak at all conditions suggests that both domains in Athe_2310 melt at similar temperatures, and that this phenomenon is intensified when it adopts its holo conformation in the presence of cognate ɑ-glucan ligands – based on increased peak narrowing.

**Figure 4:**
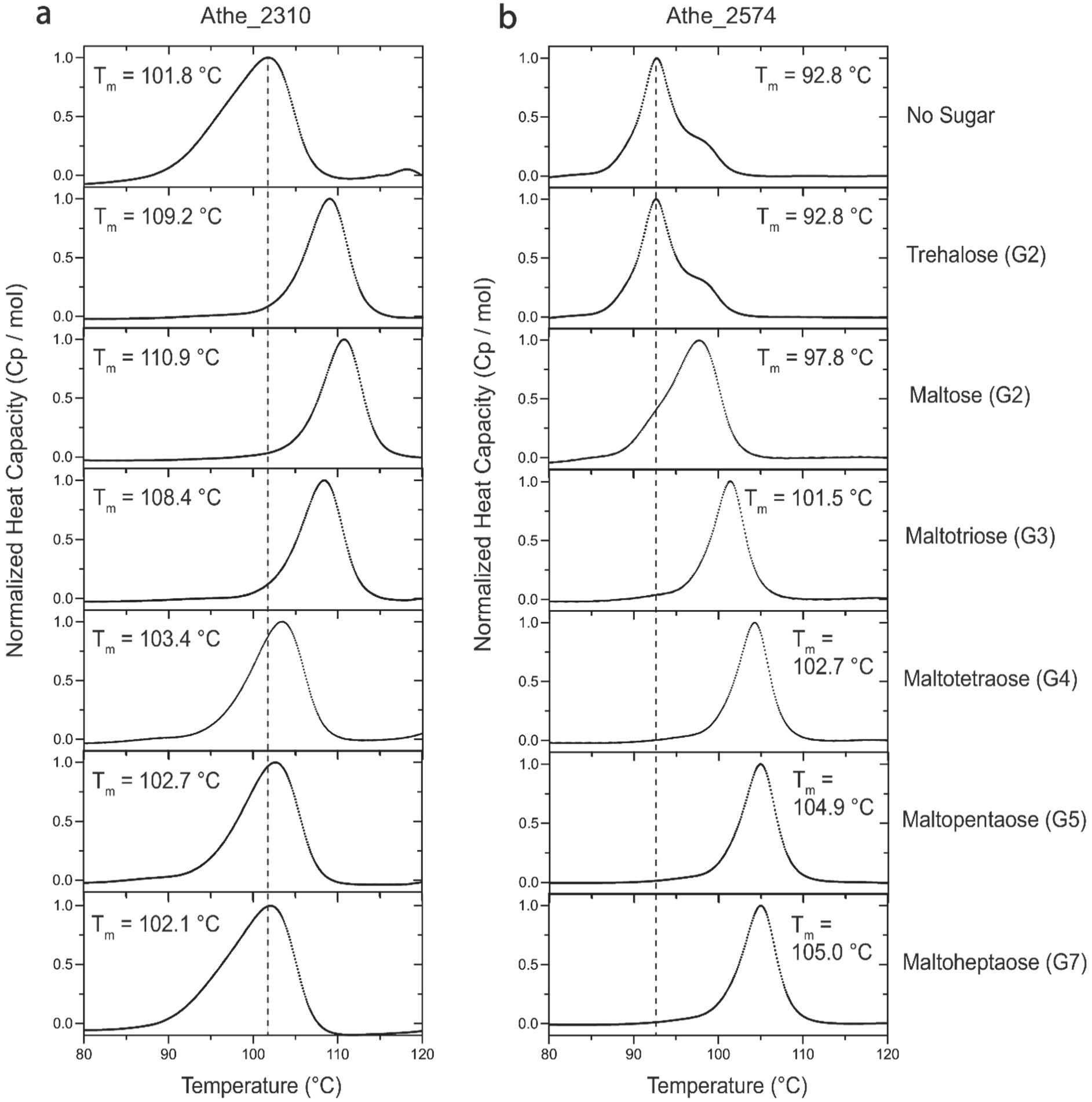
Normalized differential scanning calorimetry (DSC) screens of a) Athe_2310 and b) Athe_2574 mixed with ɑ-glucan substrates of various lengths: trehalose (G2) maltose (G2), maltotriose (G3), maltotetraose (G4), maltopentaose (G5), maltoheptaose (G7). Athe_2310 experienced its highest melting temperature shifts for shorter-length substrates (G2–G3), while Athe_2574 experienced its highest melting temperature shifts with longer maltodextrins (G4–G7). DSC screens were performed at a temperature range of 60 - 130 °C, in 50 mM HEPES and 300 mM NaCl pH 7.0 buffer.

Athe_2574 displays a distinct DSC melting profile compared to Athe_2310 (Figure 4). Firstly, our results suggest that the two domains in Athe_2574 melt at slightly different temperatures in its open, apo state: one domain melts near 90 °C while the other domain melts closer to 100 °C (Figure 4b). The inclusion of trehalose did not alter the protein melting profile, indicating an absence of binding. However, maltose appears to induce domain closure such that the melting profile of Athe_2574 is more uniform. This effect is more pronounced with increasing maltodextrin length – the presence of oligosaccharides maltotriose to maltoheptaose yields the narrowest melting curve profiles for Athe_2574. The largest melting temperature differences are also observed with these longer oligosaccharides: maltotetraose, maltopentaose, maltoheptaose, which all generated *ΔT*_m_ > 12 °C (Figure 4b). Thus, longer maltodextrins yield the greatest thermal stabilization effect on Athe_2574.

Next, ITC was performed to probe the binding interactions between our MBPs and the same palette of ɑ-glucan oligosaccharides. Isothermal titration curves and binding isotherms are shown in Figure 5, with key parameters such as dissociation constants (*K*_d_) and stoichiometry (*n*) summarized in Table 2. These data show that Athe_2310 binds to trehalose, maltose, and maltotriose with micromolar dissociation constants (Table 2). Enthalpies of binding are also exothermic in all cases, consistent with binding isotherms observed with TtMBP, PfuMBP, and *E. coli* MBP (20,21,48). As DSC experiments indicated an absence of binding between Athe_2310 and longer maltodextrins (G4–G7), ITC measurements for these mixtures were not performed. In all cases, entropic contributions to binding between Athe_2310 and each cognate ligand slightly exceed enthalpic contributions (Table 2).

**Figure 5:**
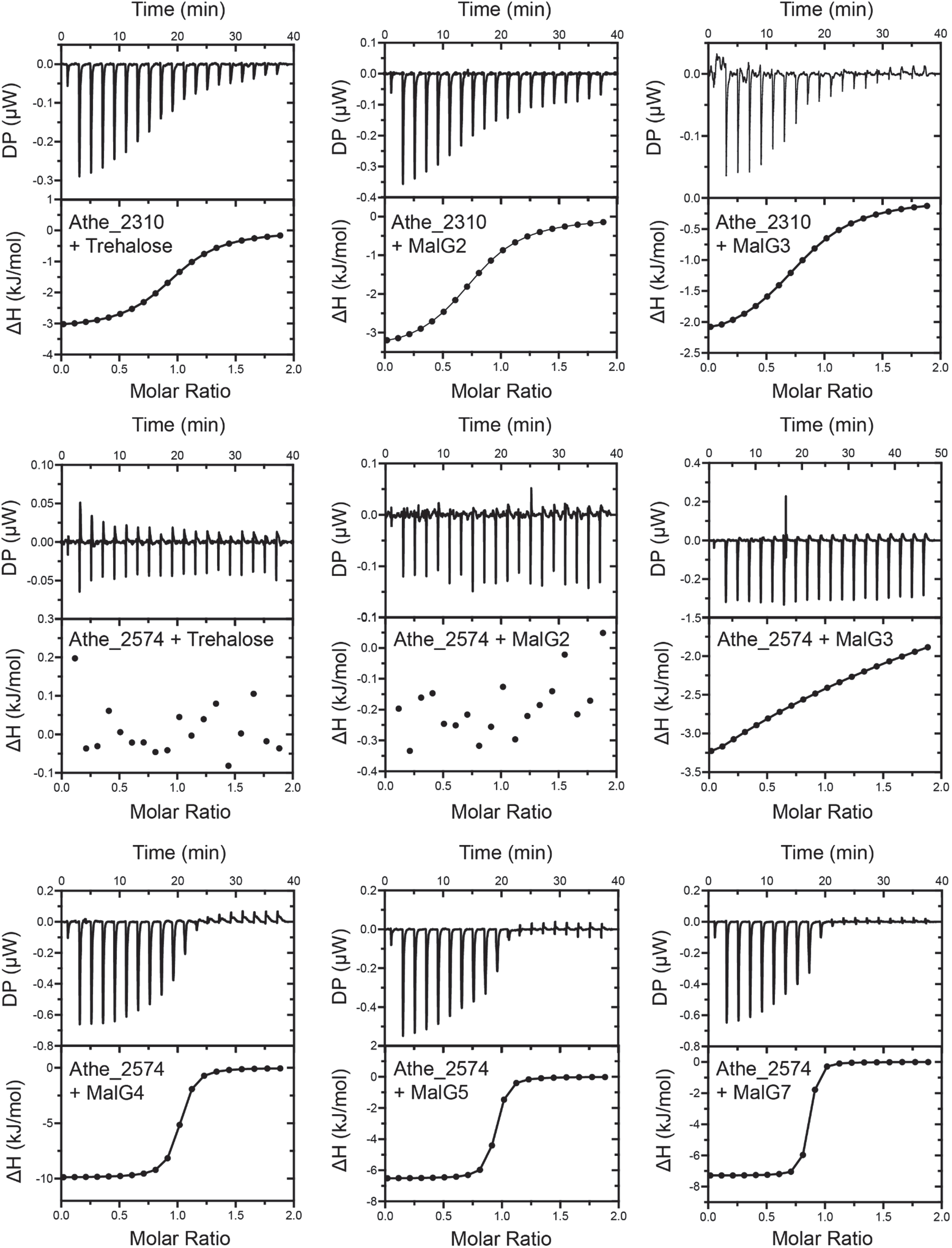
Representative isothermal titration calorimetry (ITC) screens of Athe_2310 and Athe_2574 with various ɑ-glucan substrates: trehalose, maltose (MalG2), maltotriose (MalG3), maltotetraose (MalG4), maltopentaose (MalG5), maltoheptaose (MalG7). For each AbMBP-carbohydrate mixture, both raw isothermal titration curves and integrated binding isotherms are shown.

**Table 2:** Biophysical and thermodynamic parameters of binding between MBPs and various ɑ-glucan substrates as determined by DSC and ITC. MBP-substrate combinations marked with a “-” were not tested via the ITC as DSC measurements indicated either weak or absent binding.

| Protein | Sugar | $T_m$ (°C) | $\Delta T_m$ (°C) | $n$ | $K_d$ (μM) | $K_a$ ($\times 10^5$ M<sup>-1</sup>) | $T$ (°C) | $\Delta H$ ($\frac{\text{kcal}}{\text{mol}}$) | $T\Delta S$ ($\frac{\text{kcal}}{\text{mol}}$) | $\Delta G$ ($\frac{\text{kcal}}{\text{mol}}$) |
| --- | --- | --- | --- | --- | --- | --- | --- | --- | --- | --- |
| Athe_2310 | Trehalose | 109.02 | 7.23 | 0.96<br>± 0.02 | 2.59<br>± 0.50 | 3.86 | 25 | -3.18<br>± 0.14 | -4.45 | -7.63 |
| | Maltose | 110.85 | 9.06 | 0.76<br>± 0.02 | 3.93<br>± 0.78 | 2.54 | 25 | -3.53<br>± 0.18 | -3.85 | -7.38 |
| | MalG3 | 108.42 | 6.63 | 0.79<br>± 0.06 | 5.21<br>± 2.98 | 1.92 | 25 | -2.36<br>± 0.38 | -4.85 | -7.21 |
|  | MalG4 | 103.36 | 1.57 | - | - | - | - | - | - | - |
|  | MalG5 | 102.73 | 0.95 | - | - | - | - | - | - | - |
|  | MalG7 | 102.06 | 0.26 | - | - | - | - | - | - | - |
| Athe_2574 | Trehalose | 92.69 | 0.1 | - | - | - | - | - | - | - |
|  | Maltose | 97.80 | 5.03 | - | - | - | - | - | - | - |
| | MalG3 | 101.47 | 8.72 | 1.57<br>± 85.80 | 523<br>± 27300 | 0.019 | 25 | -24.80<br>± 2410 | 20.4 | -4.48 |
| | MalG4 | 104.26 | 11.50 | 0.98<br>± 0.01 | 0.17<br>± 0.04 | 58.48 | 25 | -9.90<br>± 0.19 | 0.664 | -9.24 |
| | MalG5 | 104.93 | 12.18 | 0.90<br>± 0.00 | 0.10<br>± 0.01 | 99.01 | 25 | -6.52<br>± 0.07 | -3.03 | -9.56 |
| | MalG7 | 105.04 | 12.29 | 0.82<br>± 0.00 | 0.05<br>± 0.01 | 204.92 | 25 | -7.29<br>± 0.04 | -2.69 | -9.98 |

Athe_2574 preferentially binds longer malto-oligosaccharides, with the highest affinity for maltoheptaose (*K*_d_ = 0.049 uM at 25 °C) (Table 2). In contrast, binding affinity to maltotriose was relatively weak (*K*_d_ = 523 uM at 25 °C). The shift in binding affinity from maltotriose to maltotetraose is marked, with a 1000-fold change in *K*_d_. Distinct from Athe_2310 is the much greater relative enthalpic contributions compared to entropic contributions for ligand coordination exhibited by Athe_2574 with all sugars. And although binding between maltose and Athe_2574 is captured by DSC, enthalpy changes of binding were too small to fit meaningful *K*_d_ and *n* values. Athe_2310 exhibits dissociation constants in the single digit uM range, indicating high affinity binding consistent with observed thermophilic orthologs (33). Yet, dissociation constants an order of magnitude lower were observed for Athe_2574 and long malto-oligosaccharides (G5–G7), indicating an exceptionally high binding affinity for these substrates. Indeed, the largest thermal stabilization shifts across both MBPs and their substrates are observed between Athe_2574 and maltotetraose-maltoheptaose.

### Ligand Docking Simulations Elucidate Structural Context to Maltodextrin Binding

Results from our computational modeling recapitulated trends in binding affinity and selectivity obtained from the experiments. Predicted binding free energies for Athe_2310 indicate a preference for shorter sugars, whereas Athe_2574 exhibits more favorable binding with longer malto-oligosaccharides (Figure 6a). To further elucidate the trends in binding, we computed the electrostatic potential of the MBPs (Figure 6b) and characterized the coordination of select ligands in the respective binding pockets (Figures 6c and 6d). Both proteins have regions of negative electrostatic potential at the hinge region of the protein that may provide long-range attractive forces between the protein and the polar sugar, but Athe_2574 has a negative potential of greater magnitude (Figure 6b). Favorable binding free energies are driven by the formation of enthalpic interactions between the sugar and the protein. In energetically favorable conformations, each glucose monomer forms at least one hydrogen bond or salt bridge with the protein. A subsite is formed by a set of amino acids that interact with the same glucosyl monomer. As only two subsites are observed in Athe_2310, this suggests that disaccharides are most likely to stabilize its binding pocket (Figure 6c). On the other hand, the presence of four subsites in the lowest energy pose for maltotetraose in Athe_2574 is indicative of its relative preference for larger oligosaccharides (Figure 6d).

**Figure 6:**
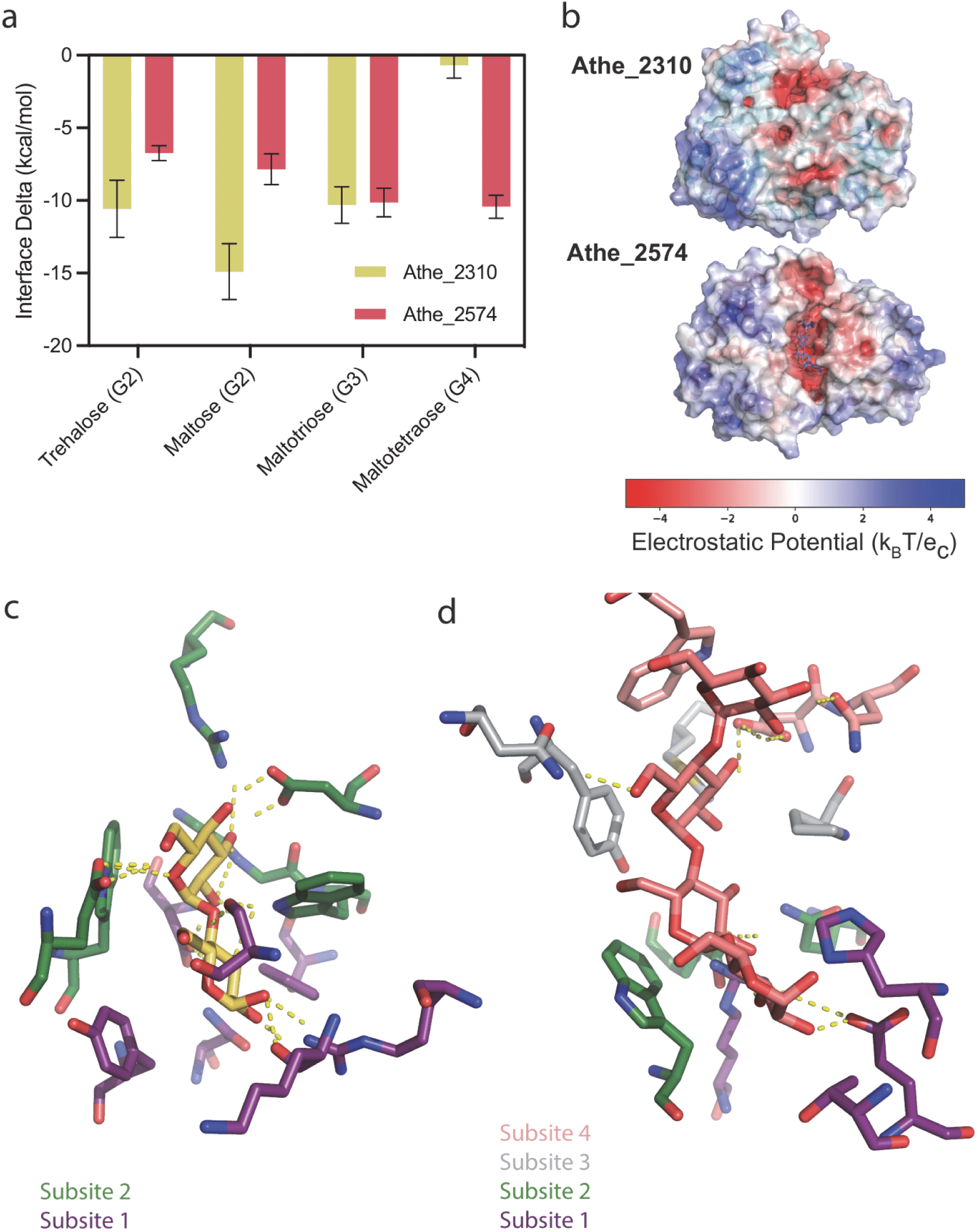
a) Interface energy deltas, corresponding to free energies of binding, for each simulated maltodextrin-binding protein and substrate combination. The rank-ordering of each protein–substrate combination is aligned with experimental results. b) Surface charge distribution, as measured by electrostatic potential, of Athe_2310 and Athe_2574 based on AlphaFold2 models. c) Coordination of cognate ligand maltose in the Athe_2310 binding pocket. d) Coordination of cognate ligand maltotetraose in the Athe_2574 binding pocket.

Substrate specificity can be linked to binding pocket subsite availability for each glucosyl monomer, and the overall orientation of the sugar. For Athe_2310, R12, N316, and conserved residues D121 and F119 comprise subsite 1, while conserved residues D69, E225, W243, R351, and W280 form subsite 2 (Figure 6c). Subsite 3 is partially concealed by a shift in the a-helix formed by residues 220–234, whereas subsite 4 is fully occluded due to the additional turn on the a-helix formed by residues 41–48, leaving only subsites 1 and 2 available for ligand binding (Figure 6c). On the other hand, Athe_2574 stabilizes the reducing end of maltotetraose via polar interactions with H8, E13 and the conserved residue R291. Polar contacts between the glycosyl and conserved residues W223, E109 and Q256 comprise subsite 2; interactions with conserved residues P61, Y147, and Y148 comprise subsite 3; and interactions with conserved residues D63 and W333 comprise subsite 4. As the pocket volume of Athe_2574 (396 Å^3^) is much greater than that of Athe_2310 (109 Å^3^), it is unsurprising that Athe_2574 possesses more available subsites. Ligand-protein interface energetic calculations and structural analysis of their respective binding pockets suggest a mechanism by which Athe_2310 and Athe_2574 discriminate between maltodextrin substrates by size.

## Discussion

Our combined biophysical and structural analysis show that Athe_2308–Athe_2310 and Athe_2574–2578 are maltodextrin transporters in *A. bescii*. Substrate-binding proteins are known to alter conformations from an open, apo state to a closed, holo state in the presence of their cognate substrates. Using differential scanning calorimetry (DSC), we probed the thermostability of Athe_2310 and Athe_2574 with and without ɑ-glucan substrates to screen for potential ligands (Figure 4). To confirm the ligands predicted by DSC, we performed isothermal titration calorimetry (ITC) to quantify the thermodynamics of oligosaccharide binding (Figure 5, Table 2). Sequence and structural alignment to adjacent thermophilic orthologs (Figures 2 and 3) provided structural insights to maltodextrin coordination, forming the basis of ligand docking simulations (Figure 6).

Without supplying a template structure, AlphaFold2 was not able to predict the corresponding number of subsites needed for maltotetraose binding in Athe_2574. As the AlphaFold training set includes holo maltose/maltodextrin binding proteins predominantly bound to maltose, the AlphaFold EvoFormer predictions give the closed maltose-bound conformation in the absence of any bound ligand (49). Supplying a template structure therefore ensures the correct number of subsites are present in the modeled structure. For the four sugars (trehalose, maltose, maltotriose, and maltotetraose) where there were crystal structures of holo orthologs, our computational rank-ordering of each sugar’s binding affinity for each receptor matched results from ITC and DSC measurements (Figure 6A).

Biophysical results from both ITC and DSC agree: preferred substrates exhibit both higher *T*_m_ differences and lower *K*_d_ constants (Table 2). Athe_2310 achieves its most thermostable conformation when bound to maltose, while Athe_2574 is most thermostable when bound to maltoheptaose. Though ITC suggests that Athe_2310 slightly prefers trehalose to maltose (*K*_d_ = 2.59 uM versus *K*_d_ = 3.93 uM at 25 °C), *ΔT*_m_ with maltose is 1.83 °C higher, suggesting that maltose is more effective at pocket stabilization. Specific yet weak interactions as indicated by a modest *ΔT*_m_ = 1.57 °C may be taking place between Athe_2310 and maltotetraose, but it is also possible that trace impurities of maltose or maltotriose are present from the manufacturer, as was similarly observed in the characterization of PfuMBP (21). It is also possible that binding affinity and precise substrate specificity as described in Table 2 may differ at higher, native temperatures (22). However, we performed our ITC measurements at room temperature due to exacerbated thermal fluctuations in the buffer that worsen signal-to-noise ratios at elevated temperatures (22,50,51).

We propose that two orthogonal ABC transporters are responsible for high-affinity maltodextrin uptake in *A. bescii*. The maltose/trehalose ABC Transporter (Athe_2308 - 2310) uptakes maltose, trehalose, and maltotriose, thereby preferring shorter maltodextrins (<G3) while the maltodextrin ABC Transporter (Athe_2574, Athe_2577 - Athe_2578) primarily uptakes longer malto-oligosaccharides (>G3).

A bioenergetic benefit to uptaking higher-order maltodextrins explains the affinity of Athe_2574 for longer oligosaccharides. A fixed cost of two ATP molecules per translocation of one sugar molecule implies that longer sugars provide higher ATP rates of return (52,53). *A. bescii* is also known to secrete glycoside hydrolases with starch-binding domains (Athe_0609 and Athe_0610) (54). And though extracellular CAZymes from the so-called Glucan Degradation Locus (GDL) in *A. bescii* (Athe_1860 - 1867) primarily degrades beta-glucans, their activities are diverse and can potentially degrade longer alpha-glucans into disaccharides and trisaccharides (55,56). Possession of an ABC transporter for smaller substrates allows *A. bescii* to mitigate risk from the extracellular availability of both long and short oligosaccharides – conferring metabolic benefit in sugar-poor environments.

Another thermophilic bacterium, *T. maritima,* is known to utilize multiple ABC maltodextrin transporters for uptaking short and long maltodextrins (32,40). Whereas *A. bescii* possesses two maltodextrin transporters, *T*. *maritima* retains three transporter operons – each utilizing its own MBP isoform: TmMBP1, TmMBP2, TmMBP3 (the substrate specificity of these MBP isoforms have been described in earlier sections) (32). Possessing genes encoding distinct maltodextrin transporters implies a larger genome size. Presumably, however, the added metabolic costs is outweighed by the benefit of additional specialized maltodextrin transporters – as opposed to a single maltodextrin transporter that can bind promiscuously between short and long oligosaccharides. For example, whereas *E. coli* MBP binds non-selectively across maltose to maltoheptaose, selective binding to only certain substrates implies greater conformational inflexibility, which leads to tighter affinity interactions (35,37). Though the lineage of these two maltodextrin transporters are not clear, it is well understood that thermophilic ABC transporters can adaptively evolve in response to available carbohydrate substrates so as to maximize organism fitness (57).

The promiscuous *msmK* ATPase (Athe_1803) likely powers both ABC maltodextrin transporters in *A. bescii* (Figure 7). No other ATPases are associated with the two maltodextrin transporters. An *A. bescii msmK-deletion* strain grew to an eight-fold lower optical density (OD680) relative to wildtype on beta-linked cellobiose (19). Promiscuous ATPases have also been identified in other gram-positive bacteria including *Streptomyces reticuli* (23,58). Still, this contrasts observations that thermophilic bacteria and archaea possess multiple copies of *malK* (39,40,59,60). And though *T. maritima* is a gram-negative bacterium with each of its maltodextrin transporters possessing a unique ATPase domain, the deletion of only one *malK* (locus tag: THMA_1301) impaired activity from all three maltodextrin transporters (40).

**Figure 7:**
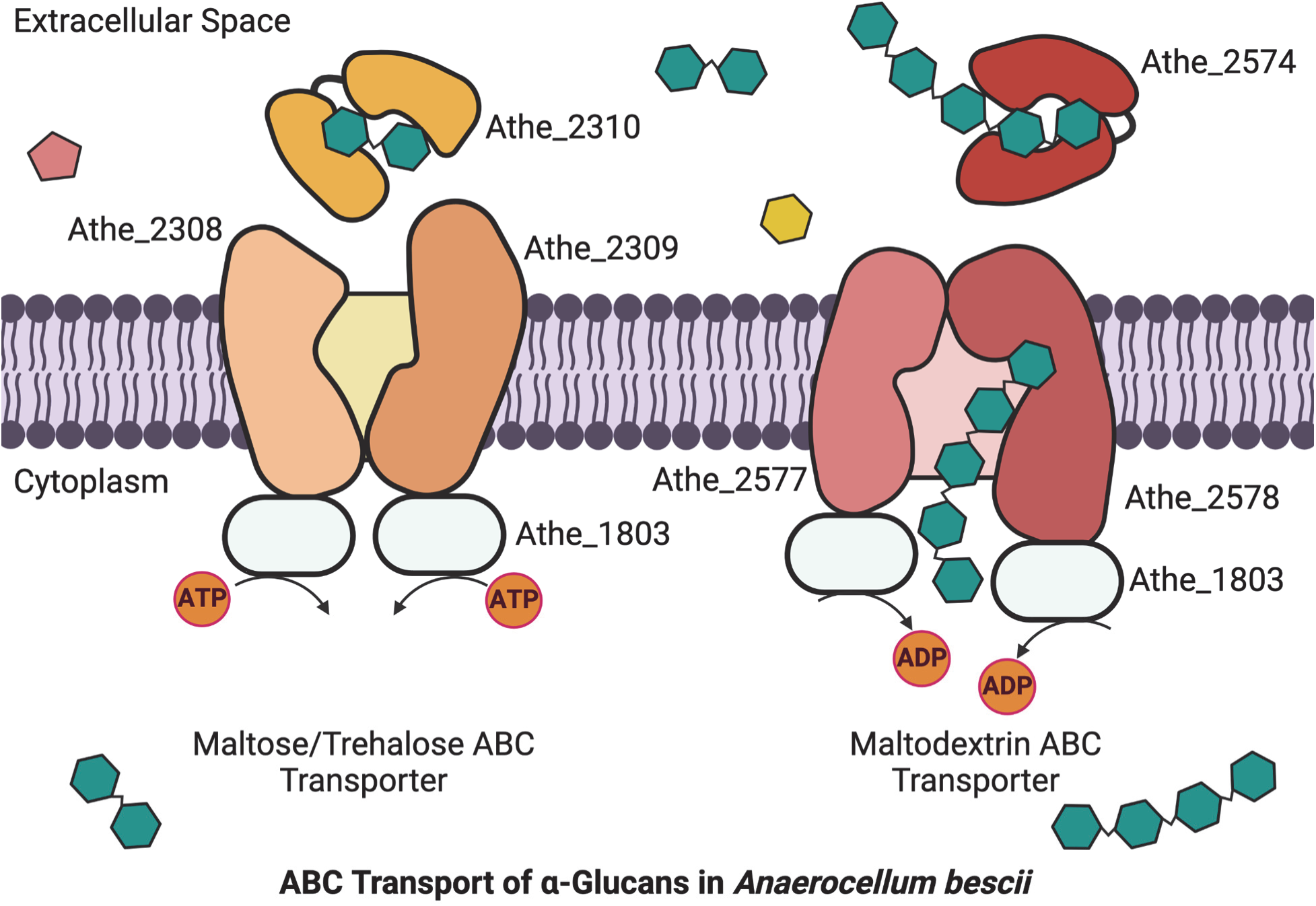
Two CUT1 ABC transporters powered by the promiscuous MsmK ATPase facilitate uptake of ɑ-glucan substrates in *A. bescii.* The maltose/trehalose ABC transporter Athe_2308-2310 preferentially uptakes shorter maltodextrins up to maltotriose, while the maltodextrin ABC transporter (Athe_2574-2578) transporters longer-chained maltodextrins such as maltotetraose and maltopentaose.

## Conclusion

In this work, we have shown that malto-oligosaccharide uptake in *A. bescii* is mediated by two, non-overlapping high-affinity ABC transporters. The ABC transporter Athe_2308 - Athe_2310 translocates short maltodextrins (up to maltotriose) across the membrane, whereas Athe_2574 - Athe_2578 uptakes long maltodextrins (up to maltoheptaose) (Figure 7). Both ABC transporters are likely powered by the promiscuous MsmK ATPase (Athe_1803) (Figure 7). The two maltodextrin-binding proteins (Athe_2310 and Athe_2574) possess similar structural features to known thermophilic orthologs, with significant conservation of binding pocket residues and subsites that likely govern substrate specificity. Athe_2310 displays similar substrate specificity to known thermophilic maltose/trehalose-binding proteins, conversely the binding of Athe_2574 to long maltodextrins G4–G7 has not been extensively studied in other thermophiles. Through ligand docking simulations on modified AlphaFold2 structures, we predicted each MBP’s subsites for maltodextrin binding that give rise to their distinct substrate specificities consistent with the behavior of known orthologs. It is likely the presence of two ABC transporters for uptake of maltodextrins of different size in *A. bescii* enables efficient scavenging in its native carbohydrate limiting environments – an adaptation that has also been observed in other thermophiles (40,57,61). This study represents a first in the identification and biophysical characterization of ABC sugar transporters in *Anaerocellum* (formerly *Caldicellulosiruptor*) species, paving the way for study of additional sugar transporters as well as their manipulation for industrial fuels and chemical production.

## Materials & Methods

### Bacteria & Growth Conditions

*E. coli* strains, NEB 5-ɑ (New England Biolabs) and BL21 (DE3) pRosetta2 (EMD Millipore), were routinely plated on Luria-Bertani medium (5 g/L of yeast extract, 10 g/L of tryptone, 10 g/L of NaCl) with 1.5% agar and 50 µg/mL kanamycin. All *E. coli* strains were routinely cultured in Luria-Bertani medium supplemented with and additional 24 g/L of yeast extract and working concentrations of kanamycin (50 µg/mL) as appropriate for selection (62). Additional 33 µg/mL chloramphenicol antibiotic was supplemented for plating and culturing *E. coli* BL21 (DE3) pRosetta2 strains. *A. bescii* DSMZ 6725 was obtained from the lab of Robert Kelly (North Carolina State University) and grown on complex media (C516) containing 0.5 g/L yeast extract, 0.5 g/L maltose substrate, and 40 µM uracil as described previously (55,63). *A. bescii* cells were harvested in the late exponential phase and pelleted by centrifugation at 5,000 x g for 20 min. Genomic DNA was purified from cell pellets using a *Quick*-DNA Miniprep kit (Zymo Research) and quantitated using a UV-Vis Spectrophotometer Nanodrop (Thermo Scientific).

### Chemicals

The following oligosaccharides were used in this study: D-Maltose Monohydrate (>98.0%, Tokyo Chemical Industry), D-Trehalose Dihydrate (>99.0%, Sigma Aldrich) D-Maltotriose (>98.0%, Carbosynth Ltd), D-Maltotetraose (>95.0%, Cayman Chemical Company)., D-Maltopentaose (>90.0%, Carbosynth Ltd), D-Maltoheptaose (>80.0%, Carbosynth Ltd).

### Substrate Binding Protein Cloning

Genes encoding Athe_2310 and Athe_2554 were cloned lacking their signal peptide (predicted by SignalP 5.0) into pRSF1-b (gift from the Kelly Lab, North Carolina State University) and pCri8a (Addgene Plasmid #61317) backbones, respectively, via Gibson Assembly (64,65). PCR was conducted using primers in Table S1 using Phusion polymerase (New England Biolabs) according to manufacturer instructions on *A. bescii* genomic DNA. Gibson Assembly was performed using NEB HiFi Assembly Mastermix according to the manufacturer instructions. Each substrate-binding protein gene contained an N-terminal hexa-histidine-tag for immobilized metal affinity chromatography (IMAC) purification. These plasmids, dubbed pHT002 and pHT003a for Athe_2310 and Athe_2554, respectively, were transformed to NEB 5-ɑ *E. coli* cells and selected on 50 ug/mL kanamycin for plasmid maintenance. Plasmids were prepared using the Zymo Research Plasmid Miniprep - Classic kit (Zymo Research) and sequenced using the Plasmidsaurus whole plasmid sequencing service. Sequence confirmed plasmid was transformed into BL21 (DE3) pRosetta2 *E. coli* (EMD Millipore) and selected on 50 ug/mL kanamycin and 33 ug/mL chloramphenicol for protein production.

### Protein Expression & Purification

Each protein expression strain was grown overnight in 1000 mL ZYM-5052 auto-induction media with 50 ug/mL kanamycin and 33 ug/mL chloramphenicol in 2.8 L shake flasks at 37 °C and 250 rpm (66). Cell culture was harvested at 4,500 x g for 20 minutes, with pellets stored in −20°C. Cell pellets were resuspended with 10 mL per g of wet cell pellet using IMAC Buffer A (20 mM sodium phosphate monobasic, 500 mM sodium chloride, pH 7.4). Resuspended pellets were lysed using an Emulsiflex-C5 High-Pressure Homogenizer (Avestin) operating on house air flow regulated at 30 psi for a target pressure of at 100,000 kPa per pass. A minimum of two full lysis cycles were performed for each lysate. Cell lysates were heat treated using a water bath set to 68°C for 30 minutes. Heat treated lysate was clarified by centrifugation at 36,000 x g for 30 minutes followed by pooling of supernatant and filtering through a 0.2 µm PES filter. This clarified cell extract was loaded into a 5 mL HisTrap HP Nickel-Sepharose (Cytiva) column, operated as per manufacturer instructions using a BioRad NGC 10 FPLC (Bio-Rad). Select Ni-NTA elution samples were run with a precast 4-20% Mini-PROTEAN® TGX Stain-Free Protein Gel (Bio-Rad), using a Precision Plus Protein Standard (Bio-Rad), against filtered lysate, sample load, and column wash samples to determine high protein purity fractions for pooling. Select purified elution fractions were pooled, and concentrated, as well as buffer-exchanged, using a 10 kDa MWCO PES filter 20 mL Spin-X Concentrator (Corning) into 50 mM HEPES, 300 mM NaCl, pH 7.0 as the final protein storage and characterization buffer. Each protein was concentrated to 50 mg/mL based on protein quantitation using the bicinchoninic acid (BCA) assay (Thermo Fisher Scientific).

### Isothermal Titration Calorimetry (ITC)

Proteins were dialyzed into the following buffer: 50 mM HEPES, 300 mM NaCl, pH 7.0 via 10 kDa MWCO PES filter 20 mL Spin-X Concentrators (Corning). Measurements were performed at 25 °C using a MicroCal PEAQ Isothermal Titration Calorimeter (Malvern Panalytical). A standard titration experiment consisted of the protein sample at 50 μM and the sugar ligand at 500 µM. Stirring was set at 750 rpm. The 280 μL sample of protein was injected with a priming aliquot of 0.2 μL of sugar ligand, followed by 19 successive injections of 2.0 μL at 120-s intervals. Integrated heat effects were determined by non-linear regression using a single site binding model (Microcal PEAQ-ITC Analysis). The fitted isotherms yield the binding dissociation constant and other thermodynamic parameters calculated using the differential form of the Gibbs Free Energy equation *ΔG*_d_ = -*RTln*(*K*_d_). The slope of the binding isotherm at the equivalence point was used to determine the dissociation constant *K*_d_. The association rate constant *K*_a_ is defined as *K*_a_ = 1/*K*_d_.

### Differential Scanning Calorimetry (DSC)

DSC measurements using a MicroCal PEAQ-DSC (Malvern Panalytical) were performed on Athe_2310 and Athe_2574 in combination with a variety of ɑ-glucans: trehalose (G2), maltose (G2), maltotriose (G3), maltotetraose (G4), maltopentaose (G5), and maltoheptaose (G7). The scan rate was set to 180 °C / min, with a max temperature of 130 °C. Protein was mixed with each sugar, yielding final concentrations of 2.5 mg/mL of protein and 5 mM of sugar. 50 mM HEPES, 300 mM NaCl, pH 7.0 was used as the reference buffer. The heat capacity *C*_p_ was plotted against temperature and then normalized by the respective run’s maximum *C*_p_ value (*C*_p,max_). We defined the melting temperature *T*_m_ as the temperature corresponding to *C*_p,max_ = *C*_p_(*T* = *T*_m_). The melting temperature difference *ΔT*_m_ = |*T*_m,holo_ - *T*_m,apo_| is defined as the absolute difference in melting temperature between the melting temperature of a protein-sugar mix and the same protein with no sugar present.

### Bioinformatics

Through PDB sequence similarity search, six thermophilic maltose-binding protein orthologs with solved crystal structures were identified using BLAST (67). These structures were from *Thermococcus litoralis (PDB: 1EU8), Thermotoga maritima (PDB: 6DTQ, PDB: 6DTS, PDB: 6DTU), Thermus thermophilus (PDB: 6J9W),* and *Pyroccocus furiosus (PDB: 1ELJ)* (20,21,32,33). A local BLAST search for each Athe_2310 and Athe_2574 as query protein sequences was performed to determine their closeness to these aforementioned orthologs, which showed that Athe_2310 and Athe_2574 are similar to a non-overlapping subset of thermophilic orthologs. Structure-based sequence alignment for each MBP and their orthologs was performed using PROMALS3D and ESPRIT visualization software to identify similarities in secondary and tertiary protein structures (43,44). PDB structure 6J9W was used to assign secondary structure to Athe_2310 and PDB structure 1ELJ was used to assign secondary structure to Athe_2574.

### Ligand Docking Models

Initial models were generated using ColabFold v1.5.5 with AlphaFold2 parameters (41,68). For Athe_2310, PDB structures 6J9W, 1EU8, and 6DTQ were used as templates. For Athe_2574, PDB structures 1ELJ, 6DTS, and 6DTU were used as templates to AlphaFold2. AlphaFold structures were then minimized according to the Rosetta *relax* protocol using the “ref2015” score function. Protein-ligand docking with RosettaLigand was performed using the ROSIE server (69,70). Sugars were initially posed in the pocket from homology to the corresponding crystal structure. 200 conformers of each ligand were generated using BCL (71). 200 docked structures for each sugar in each protein were output, and interface energies were calculated from the ten lowest energy structures. Pocket volumes were measured by the Fpocket webserver (72). Surface electrostatic potentials are calculated using APBS (Adaptive Poisson-Boltzmann Solver) (73). All protein structures were visualized using PyMOL (74).

## Acknowledgements

This work was supported by a Roberto Rocca Graduate Fellowship from Techint Group to H.T.; by the High Meadows Environmental Institute at Princeton University through the generous support of the William Clay Ford, Jr ‘79 and Lisa Vanderzee Ford ‘82 Graduate Fellowship Fund to H.T.; by a National Science Foundation Graduate Research Fellowship DGE-2039656 to V.J.; by a Gordon Wu Fellowship to V.J.; by startup funds from the Department of Chemical and Biological Engineering and the Omenn-Darling Bioengineering Institute at Princeton University to J.A.J.; by seed funds from the School of Engineering and Applied Science and startup funds from the Department of Chemical and Biological Engineering at Princeton University to J.M.C.. We acknowledge Venu Gopal Vandavasi, Director of the Princeton Biophysics Core, for instrument training and advice on experimental design. We also acknowledge Tom Silhavy and Ned S. Wingreen for helpful discussions. Finally, we acknowledge the Prud’homme Lab of Princeton University for their generous gifts of maltodextrin sugars and homogenizer.

## Conflict of Interest

The authors declare that they have no competing interests

## Notes

### Competing Interest Statement

The authors have declared no competing interest.

