## Supplementary Information for "Maltodextrin Transport in the Extremely Thermophilic, Lignocellulose Degrading Bacterium *Anaerocellum bescii (f. Caldicellulosiruptor bescii)*"

Author Affiliations:

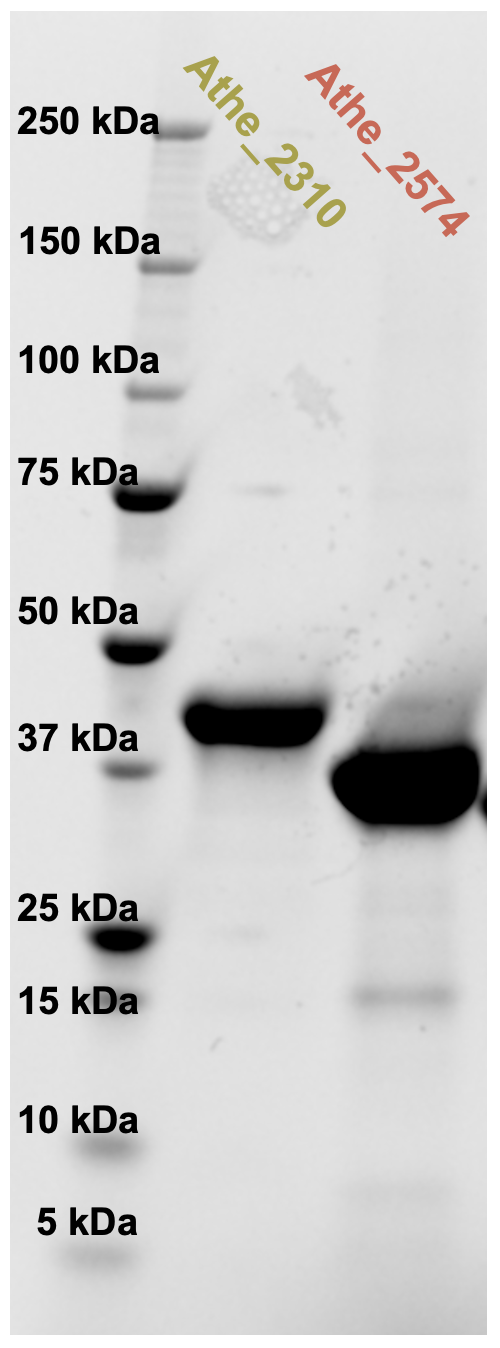

**Figure S1**: Samples of heterologously purified Athe_2310 and Athe_2574 ran on SDS-PAGE. Although Athe_2574 with its signal peptide removed is expected to weigh 40.6 kDa, it ran faster than what would be expected.

**Table S1**: Table of primers used in this study.

| Primer | DNA Sequence (5’ – 3’) | Application |
| --- | --- | --- |
| HT001_F | GCTGCCACCGCTGAGCAATAACTAG | pRGB001 Vector.FOR (backbone linearization) |
| HT002_F | cttgtcgtcgtcatccacgtgatg | pRGB001 Vector.REV (backbone linearization) |
| HT005 | CACGTGGATGACGACGACAAGGCTTCTAAAAAACAGGTCACAATTACTTATGTTCG | pHT002 Fragment FOR  (Insert amplification from *A. bescii* genomic DNA) |
| HT006 | TTGCTCAGCGGTGGCAGCTTACTTTGAAGTTTTTACAACTTCTTTTAATTTCTTGTCC | pHT002 Fragment REV  (Insert amplification from *A. bescii* genomic DNA) |
| HT134 | tgagatccggctgctaacaaag | pCri8a Vector.FOR (backbone linearization) |
| HT135 | catggcgccctggaagtaaag | pCri8a Vector.REV (backbone linearization) |
| HT136 | tttacttccagggcgccatggttACATCCAAAAAACAGCTTGTTGTC | pHT003a Fragment.FOR Athe_2574  (Insert amplification from *A. bescii* genomic DNA) |
| HT137 | ctttgttagcagccggatctcaTTATTGCATCTGAGCAATACCTTGCTTG | pHT003a Fragment.REV Athe_2574  (Insert amplification from *A. bescii* genomic DNA) |

**Table S2**: Sequences of all proteins analyzed in this study. Residues comprising the signal peptide sequence are marked in red.

| Primer | Amino Acid Sequence |
| --- | --- |
| Athe_2310  (native with signal peptide) | MKRFIAVMVLIAFSVGLFLAFGPANSNAASKKQVTITYVRGKDETHATEKIIKEFMKKNPDINVIYKENPSDTGQNHDQLVTVLSAGGSDIDVFDMDVIWPAEFAQAGYTLPLDRFIKRDKTNLNDYIKGTIDAARFKGQMWAFPRFIDAGLLYYRKDIVPQNELPKTWDDLIKVAKKYKGKNGTKYGFLMQAKQYEGLVCDAIEYIASYGGKVVDESGNIVVNNQGTIDGLNMMRKVITSGIVPPNINTFTEVETHTAFINGLSVFARNWPYMWAMINSPQSKVRGKVGILPLPKGSKGSAACLGGWMVGINKFSKNPEASWRLLKFLVQKEGQKLMAIYNGNVPVYKPLFNDKDVIKANPLIGDKKFIEAILAAVPRPVSPIYPKISDVMQIELSNIVNGKKDVKTAVADMDKKLKEVVKTSK |
| Athe_2574  (native with signal peptide) | MKNLKRILTVALIITFAVVALIPLSGVFATSKKQLVVWSHLTQDEVKALQPIADKWGKENGYTVKVITDQGSFQSFQTAAMSGKGPDIMFGIPHDNLGAFWKAKLLEAVPANLIDKKNFVSTALDACSFEGKLYALPIAMETYALFYNTSKVKEAPKTMSQLITLAKKYGFMYDVNNFYFSFAFIAQNGGYVFKNKGGSLDPNDIGLATNGAIKGLSLIRDFVQTYKFMPKDIKGDIAKGNFQNQKIAFYISGPWDVQDFIKAKVPFAVAPLPKTDDGKPTPSFVGVQAAFVSAKSKNKDAAFKLMKYLVENSALTLFKVGHRIPVLNKVLTSSEVKADKIMSAFAEQAKVGIPMPNIPEMSAVWPVANNALSLITTGKATPKQAADAMVKQIKQGIAQMQ |
| Athe_2310  (cloned in *E. coli*) | MAHHHHHHVDDDDKASKKQVTITYVRGKDETHATEKIIKEFMKKNPDINVIYKENPSDTGQNHDQLVTVLSAGGSDIDVFDMDVIWPAEFAQAGYTLPLDRFIKRDKTNLNDYIKGTIDAARFKGQMWAFPRFIDAGLLYYRKDIVPQNELPKTWDDLIKVAKKYKGKNGTKYGFLMQAKQYEGLVCDAIEYIASYGGKVVDESGNIVVNNQGTIDGLNMMRKVITSGIVPPNINTFTEVETHTAFINGLSVFARNWPYMWAMINSPQSKVRGKVGILPLPKGSKGSAACLGGWMVGINKFSKNPEASWRLLKFLVQKEGQKLMAIYNGNVPVYKPLFNDKDVIKANPLIGDKKFIEAILAAVPRPVSPIYPKISDVMQIELSNIVNGKKDVKTAVADMDKKLKEVVKTSK |
| Athe_2574  (cloned in *E. coli*) | MGSSHHHHHHSSGENLYFQGAMVTSKKQLVVWSHLTQDEVKALQPIADKWGKENGYTVKVITDQGSFQSFQTAAMSGKGPDIMFGIPHDNLGAFWKAKLLEAVPANLIDKKNFVSTALDACSFEGKLYALPIAMETYALFYNTSKVKEAPKTMSQLITLAKKYGFMYDVNNFYFSFAFIAQNGGYVFKNKGGSLDPNDIGLATNGAIKGLSLIRDFVQTYKFMPKDIKGDIAKGNFQNQKIAFYISGPWDVQDFIKAKVPFAVAPLPKTDDGKPTPSFVGVQAAFVSAKSKNKDAAFKLMKYLVENSALTLFKVGHRIPVLNKVLTSSEVKADKIMSAFAEQAKVGIPMPNIPEMSAVWPVANNALSLITTGKATPKQAADAMVKQIKQGIAQMQ |
| TmMBP3 (PDB: 6DTQ) | MAVKITMTSGGVGKELEVLKKQLEMFHQQYPDIEVEIIPMPDSSTERHDLYVTYFAAGETDPDVLMLDVIWPAEFAPFLEDLTADKDYFELGEFLPGTVMSVTVNGRIVAVPWFTDAGLLYYRKDLLEKYGYDHAPRTWDELVEMAKKISQAEGIHGFVWQGARYEGLVCDFLEYLWSFGGDVLDESGKVVIDSPEAVAALQFMVDLIYKHKVTPEGVTTYMEEDARRIFQNGEAVFMRNWPYAWSLVNSDESPIKGKVGVAPLPMGPGGRRAATLGGWVLGINKFSSPEEKEAAKKLIKFLTSYDQQLYKAINAGQNPTRKAVYKDPKLKEAAPFMVELLGVFINALPRPRVANYTEVSDVIQRYVHAALTRQTTSEDAIKNIAKELKFLLGQHHHHHH |
| TMBP (PDB: 1EU8) | IEEGKIVFAVGGAPNEIEYWKGVIAEFEKKYPGVTVELKRQATDTEQRRLDLVNALRGKSSDPDVFLMDVAWLGQFIASGWLEPLDDYVQKDNYDLSVFFQSVINLADKQGGKLYALPVYIDAGLLYYRKDLLEKYGYSKPPETWQELVEMAQKIQSGERETNPNFWGFVWQGKQYEGLVCDFVEYVYSNGGSLGEFKDGKWVPTLNKPENVEALQFMVDLIHKYKISPPNTYTEMTEEPVRLMFQQGNAAFERNWPYAWGLHNADDSPVKGKVGVAPLPHFPGHKSAATLGGWHIGISKYSDNKALAWEFVKFVESYSVQKGFAMNLGWNPGRVDVYDDPAVVSKSPHLKELRAVFENAVPRPIVPYYPQLSEIIQKYVNSALAGKISPQEALDKAQKEAEELVKQYS |
| TtMBP (PDB: 6J9W) | QSGPVIRVAGDSTAVGEGGRWMKEMVEAWGKKTGTRVEYIDSPADTNDRLALYQQYWAARSPDVDVYMIDVIWPGIVAPHALDLKPYLTEAELKEFFPRIVQNNTIRGKLTSLPFFTDAGILYYRKDLLEKYGYTSPPRTWNELEQMAERVMEGERRAGNRDFWGFVFQGKPYEGLTCDALEWIYSHGGGRIVEPDGTISVNNGRAALALNRAHGWVGRIAPQGVTSYAEEEARNVWQQGNSLFMRNWPYAYALGQAEGSPIRGKFGVTVLPKASADAPNAATLGGWQLMVSAYSRYPKEAVDLVKYLASYEVQKDNAVRLSRLPTRPALYTDRDVLARNPWFRDLLPVFQNAVSRPSDVAGARYNQVSEAIWTEVHSVLTGRKKGEQAVRDLEARIRRILRHHHHHH |
| PfuMBP (PDB: 1ELJ) | MKIEEGKVVIWHAMQPNELEVFQSLAEEYMALCPEVEIVFEQKPNLEDALKAAIPTGQGPDLFIWAHDWIGKFAEAGLLEPIDEYVTEDLLNEFAPMAQDAMQYKGHYYALPFAAETVAIIYNKEMVSEPPKTFDEMKAIMEKYYDPANEKYGIAWPINAYFISAIAQAFGGYYFDDKTEQPGLDKPETIEGFKFFFTEIWPYMAPTGDYNTQQSIFLEGRAPMMVNGPWSINDVKKAGINFGVVPLPPIIKDGKEYWPRPYGGVKLIYFAAGIKNKDAAWKFAKWLTTSEESIKTLALELGYIPVLTKVLDDPEIKNDPVIYGFGQAVQHAYLMPKSPKMSAVWGGVDGAINEILQDPQNADIEGILKKYQQEILNNMQG |
| TmMBP1 (PDB: 6DTU) | MQPKLTIWCSEKQVDILQKLGEEFKAKYGVEVEVQYVNFQDIKSKFLTAAPEGQGADIIVGAHDWVGELAVNGLIEPIPNFSDLKNFYETALNAFSYGGKLYGIPYAMEAIALIYNKDYVPEPPKTMDELIEIAKQIDEEFGGEVRGFITSAAEFYYIAPFIFGYGGYVFKQTEKGLDVNDIGLANEGAIKGVKLLKRLVDEGILDPSDNYQIMDSMFREGQAAMIINGPWAIKAYKDAGIDYGVAPIPDLEPGVPARPFVGVQGFMVNAKSPNKLLAIEFLTSFIAKKETMYRIYLGDPRLPSRKDVLELVKDNPDVVGFTLSAANGIPMPNVPQMAAVWAAMNDALNLVVNGKATVEEALKNAVERIKAQIQS |
| TmMBP2 (PDB: 6DTS) | MQTKLTIWCSEKQVDILQKLGEEFKAKYGIPVEVQYVDFGSIKSKFLTAAPQGQGADIIVGAHDWVGELAVNGLIEPIPNFSDLKNFYDTALKAFSYGGKLYGVPYAMEAVALIYNKDYVDSVPKTMDELIEKAKQIDEEYGGEVRGFIYDVANFYFSAPFILGYGGYVFKETPQGLDVTDIGLANEGAVKGAKLIKRMIDEGVLTPGDNYGTMDSMFKEGLAAMIINGLWAIKSYKDAGINYGVAPIPELEPGVPAKPFVGVQGFMINAKSPNKVIAMEFLTNFIARKETMYKIYLADPRLPARKDVLELVKDNPDVVAFTQSASMGTPMPNVPEMAPVWSAMGDALSIIINGQASVEDALKEAVEKIKAQIEKGSHHHHHH |
